# Comprehensive Assessment of a Fasting-Mimicking Diet: Salivary Metabolite Profiles, Weight Loss, and Biomarker Analysis

**DOI:** 10.1101/2024.12.29.630652

**Authors:** Kevin Sebastiaan Hof, Christopher Tadhg Wall

**Affiliations:** Maven Health GmbH

## Abstract

Fasting-mimicking diets (FMDs) have shown to result in various health benefits, including body weight reduction, body fat reduction, decreased blood pressure and longevity promoting effects through enhanced ketogenesis and reduced glycolysis. The direct impact of FMDs on metabolite fluctuations, however, remains elusive. Here, we explore the effects of FMDs on the human salivary metabolic profiles and weight in a single-arm case study. Specifically, we present a bio-energetic framework to estimate fat loss as a component of the sustained weight loss, the effects on metabolite quantities involved in energetic reprogramming and the effects on the abundance of metabolites originating from the gut microbiome. These findings highlight the importance of metabolic health tracking to further our understanding of the effects of fasting, personalized nutrition and effective preventive health measures.

**In brief:** Hof & Wall demonstrate that fasting-mimicking diets have short-term effects on the energy production, energetic reprogramming and gut microbiome output in humans by use of saliva metabolomics and nuclear magnetic resonance (NMR).

**Highlights:**

- 45% of total sustained weight loss could be attributed to fat loss as calculated by bio-energetic calculation framework
- Fasting-mimicking diets increase L-glutamine, L-glutamate and succinic acid indicating immediate effects on energy metabolism
- Fasting-mimicking diets increase short-chain fatty acids i.e. acetate, butyrate and propionate. Other gut microbiome metabolites are increased as well i.e. acetoin, fucose and ethanol.

## Introduction

Fasting is the voluntary abstinence from food (and drink) for a specified period and has been part of human culture throughout its history for cosmetic, religious, medical or unvoluntary (i.e. famine, hardship etc.) reasons^1^. Several types of diets exist with some focusing on eating at specific times of day with time-restricted feeding or the 16:8 method where one fasts for 16 hours and eats within an 8-hour period, called intermittent fasting. In the class of extended fasting diets (24h+) there are water fasts (where only water is consumed) and modified fasting regimes. In the modified fasting regime often very few calories and specific foods are consumed for an extended period (24h+).

Fasting-mimicking diets (FMDs) are such a modified fasting regime and have been shown to induce the health benefits of extended fasting, without the need for prolonged food deprivation^2^. FMDs are designed to lead to the same metabolic health benefits as fasting, including reduced body weight, reduced body fat, decreased blood pressure and decreased insulin-like growth factor-I (IGF-I) when compared to a normal diet^3^. A downregulation of the IGF-I pathway has been linked to improved longevity in the model organisms *C. elegans*^4^ and mice^5^. Modulation of the IGF-I pathway was shown to extend lifespan, and low concentrations of IGF-I and insulin are linked with protection against many aging-associated pathologies, including cancer and diabetes. FMDs have also been shown to improve microbial function and reduce inflammation and bowel pathology^6^. FMDs thus show a wide range of health benefits making them a compelling choice for individuals who want to reap the health benefits in an optimal manner without committing to traditional fasting protocols.

Slight adjustments in diet or lifestyle often lead to subtle metabolic changes that can be detected after a period of several months. These metabolic changes are strongly dependent on the severity of the lifestyle adjustment. In the case of severe dietary adaptations such as FMDs, the changes in human metabolism are much more rapid, with changes measurable in a matter of days^7^. Tracking of the metabolic status in the body before, during, and after lifestyle interventions like the FMDs is key to ensure their optimal effect. However, bio-fluids that are collected invasively, like blood, are a hurdle for frequent testing. In contrast, saliva is an abundant bio-fluid that can be collected non-invasively and repeatedly. Saliva metabolites reflect systemic metabolic processes and offer an accessible and practical option for monitoring the changes of the metabolic status during dietary interventions ^8^ such as FMDs. Saliva testing lends itself exceptionally well to frequent and repeated testing situations due to its non-invasive collection and ease of use. Therefore, saliva biomarker monitoring offers a valuable contribution to understanding the metabolic effects of FMDs on human physiology.

This study focused on the tracking of salivary metabolites as indicators of the metabolic changes in response to an FMD. In this white paper, we present the results of a comprehensive study on the metabolic profiles over time (pre, during and post diet) from participants undergoing the “Eat By Alex” FMD (‘reSET’ diet). By measuring key metabolites over time, we gained a deeper insight into human energy metabolism and evaluated the effects of FMDs in a short time span. Concretely, we took a closer look at the effects on the ketone body acetone and the energetic reprogramming through amino acids and succinic acid. Additional findings are expected on the response of the gut microbiota and anti-inflammatory effects by the measurements and tracking of short-chain fatty acids and other metabolites related to the gut microbiome.

## Materials & Methods

### Study design

This study was designed as single-arm trial involving 6 participants who followed a custom FMD (named the ‘reSET’ diet) for a duration of 5 days. All participants declared to adhere to the same dietary exact defined intervention. A comparison is drawn between the zero measurement (day 0, before the start of the diet) and day 5, after completion of the program, in all participants. In 2 participants the saliva samples were collected daily throughout the program, and at day 6, i.e. one day after the participant returns to their standard diet (also called refeeding).

### Participants

6 healthy adults joined this study as part of a product collaboration. These participants volunteered from the Eat By Alex (4) and Maven Health (2) teams. Inclusion criteria were to have a normal or higher BMI, while exclusion criteria were a BMI class of underweight or below, chronic illnesses, pregnancy, and eating disorders. All participants provided informed consent for the anonymized sharing of the results.

### Dietary intervention

Participants followed a structured FMD (reSET) for 5 days, consisting of low-calorie, plant-based meals designed to induce fasting-like effects. The macro- and micronutrient composition was carefully selected to aid this purpose. Compared to a normal diet, as defined by the Dietary Reference Intakes guidelines below, the reSET diet is relatively low in carbohydrates and proteins and high in (healthy) fats. All daily macronutrient intakes of the reSET diet can be seen in Table 1 and Table 2.

**Table 1.**
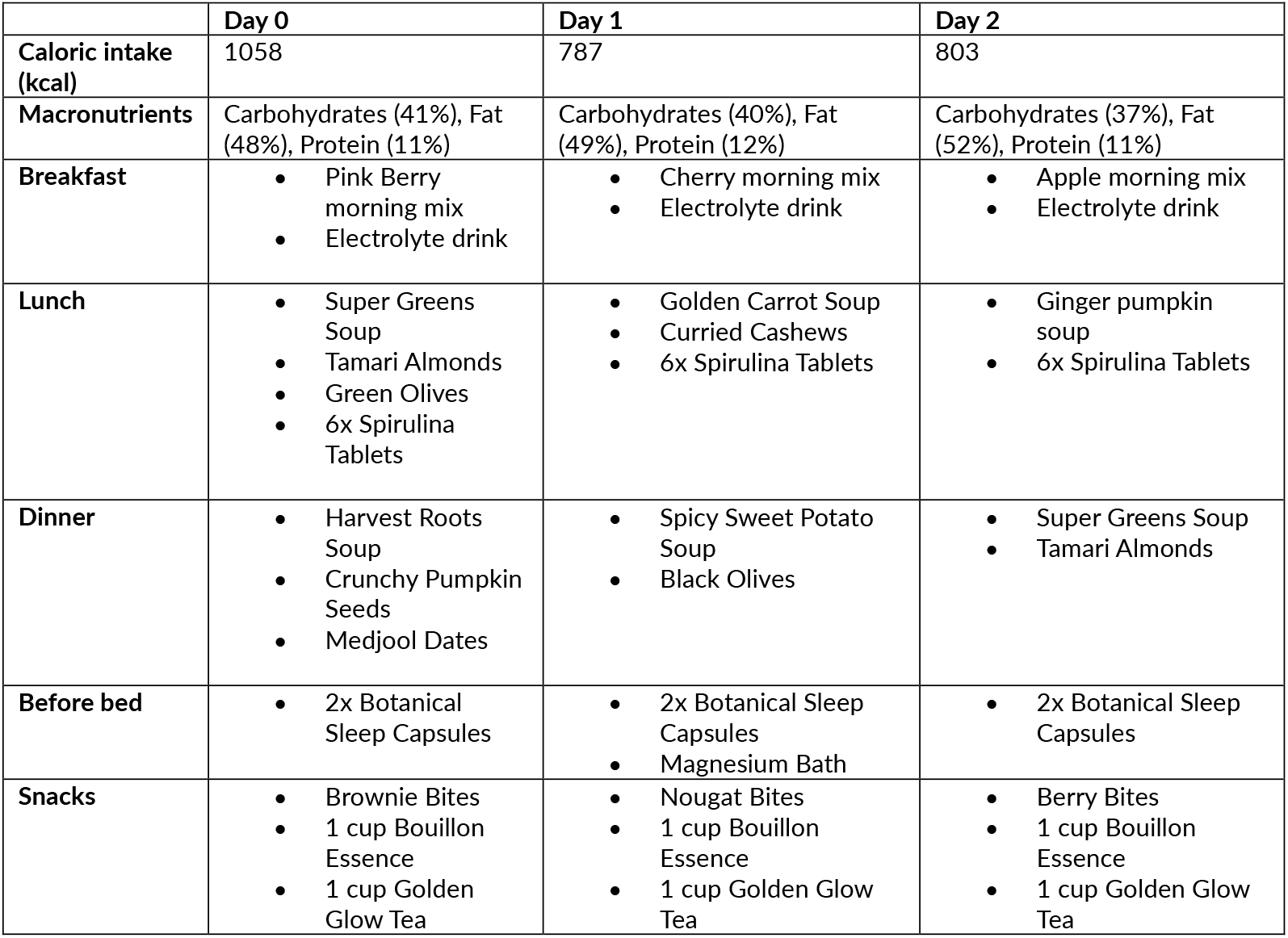
Daily overview of calories, macronutrients and all food components; Day 0-2.

|  | Day 0 | Day 1 | Day 2 |
| --- | --- | --- | --- |
| **Caloric intake (kcal)** | 1058 | 787 | 803 |
| **Macronutrients** | Carbohydrates (41%), Fat (48%), Protein (11%) | Carbohydrates (40%), Fat (49%), Protein (12%) | Carbohydrates (37%), Fat (52%), Protein (11%) |
| **Breakfast** | • Pink Berry morning mix<br>• Electrolyte drink | • Cherry morning mix<br>• Electrolyte drink | • Apple morning mix<br>• Electrolyte drink |
| **Lunch** | • Super Greens Soup<br>• Tamari Almonds<br>• Green Olives<br>• 6x Spirulina Tablets | • Golden Carrot Soup<br>• Curried Cashews<br>• 6x Spirulina Tablets | • Ginger pumpkin soup<br>• 6x Spirulina Tablets |
| **Dinner** | • Harvest Roots Soup<br>• Crunchy Pumpkin Seeds<br>• Medjool Dates | • Spicy Sweet Potato Soup<br>• Black Olives | • Super Greens Soup<br>• Tamari Almonds |
| **Before bed** | • 2x Botanical Sleep Capsules | • 2x Botanical Sleep Capsules<br>• Magnesium Bath | • 2x Botanical Sleep Capsules |
| **Snacks** | • Brownie Bites<br>• 1 cup Bouillon Essence<br>• 1 cup Golden Glow Tea | • Nougat Bites<br>• 1 cup Bouillon Essence<br>• 1 cup Golden Glow Tea | • Berry Bites<br>• 1 cup Bouillon Essence<br>• 1 cup Golden Glow Tea |

**Table 2.** Daily overview of calories, macronutrients and all food components; Day 3-4.

|  | Day 3 | Day 4 |
| --- | --- | --- |
| **Caloric intake Total (kcal)** | 790 | 762 |
| **Macronutrients** | Carbohydrates (38%), Fat (50%), Protein (12%) | Carbohydrates (37%), Fat (52%), Protein (11%) |
| **Breakfast** | • Pink Berry morning mix<br>• Electrolyte drink | • Cherry morning mix<br>• Electrolyte drink |
| **Lunch** | • Super Greens Soup<br>• Tamari Almonds<br>• Green Olives<br>• 6x Spirulina Tablets | • Golden Carrot Soup<br>• Curried Cashews<br>• 6x Spirulina Tablets |
| **Dinner** | • Harvest Roots Soup<br>• Crunchy Pumpkin Seeds<br>• Medjool Dates | • Spicy Sweet Potato Soup<br>• Black Olives |
| **Before bed** | • 2x Botanical Sleep Capsules | • 2x Botanical Sleep Capsules<br>• Magnesium Bath |
| **Snacks** | • Brownie Bites<br>• 1 cup Bouillon Essence<br>• 1 cup Golden Glow Tea | • Nougat Bites<br>• 1 cup Bouillon Essence<br>• 1 cup Golden Glow Tea |

**Table 3.** Participant parameter overview.

| Parameters | Days of collection | Mode of collection |
| --- | --- | --- |
| Age (years) | Day 0 | Self-reported |
| Sex (male/female) | Day 0 | Self-reported |
| Height (cm) | Day 0 | Self-reported |
| Number of cigarettes smoked in last 30 days | Day 0 | Self-reported |
| Weight (kg) | Day 0 – Day 8 (up to 3 days after the program had finished) | Self-reported (by each participants own weighing scale) |

The Dietary Reference Intakes (DRIs) from the Food and Nutrition Board of the Institutes of Medicine (IOM; Governmental guidelines in the USA) are defined as follows:

- Carbohydrate (45%-65%)
- Fat (20%-35%; with a limitation on saturated and trans fats)
- Protein (10%-35%)

During the diet there were three daily scheduled meals (breakfast, lunch, and dinner) and one freely administered meal (snacks). The listed snacks were eaten at the participants’ own convenience throughout the day.

### Sample & data collection

Saliva sample collection took place at the beginning and end of the diet for all participants:

- Day 0 before breakfast (non-fasting; start)
- Day 5 before breakfast (fasting; finish)

Additional saliva collection moments for the 2 participants with extensive testing:

- Day 1 before breakfast (fasting)
- Day 2 before breakfast (fasting)
- Day 3 before breakfast (fasting)
- Day 4 before breakfast (fasting)
- Day 6 before breakfast (non-fasting; +1 day after finish)

Saliva samples were collected in the morning at the earliest convenience of each participant. This is the moment of waking up before breakfast in a fasted state, not having eaten anything for a 2-hour period (pre-breakfast, brushed teeth etc.).

Saliva was collected with the Maven Health Saliva Collection Kit, which is comprised of the following items:

- 5 mL tube
- A screw cap
- A funnel
- A sticker with a unique lot number
- A biosafety collection bag

### Additional parameters

The saliva collection was performed as per Maven Health kit instructions. Briefly: participants were instructed to refrain from consuming food, chewing gum, drinking beverages other than water, brushing teeth, using mouthwash, or smoking for two hours prior to sample collection to minimize potential contamination. Immediately before collection, participants rinsed their mouths thoroughly three times using moderate amounts of tap water, allowing for the removal of residual contaminants and optimizing the collection environment. Participants were then instructed to sit comfortably and hold the tube upright while positioning the provided funnel securely in front of their teeth. Participants were asked to tilt their heads slightly forward, allowing saliva to accumulate naturally in the tube, refraining from spitting or otherwise stimulating saliva production. Collection continued until the 1 mL saliva was collected. The sealed and labeled tubes were placed in a provided safety bag for secure storage. Samples were stored at −20°C until retrieval for analysis.

### Spectra acquisition

1H-NMR spectroscopy saliva samples were rapidly thawed on ice and vortexed for 1 minute to ensure homogeneity. The samples were then centrifuged at 10,000 x g for 1 hour at 4°C to remove particulates. Following centrifugation, the cleared supernatant was transferred to a clean microcentrifuge tube. A volume of 540 µL of the supernatant was combined with 60 µL of NMR sample buffer, which contained 700 mM phosphate buffer (pH 7), 500 µM TMSP (trimethylsilyl propionate) as an internal standard, 0.5% sodium azide, and deuterium oxide (D2O) to facilitate locking during NMR acquisition. The final mixture (600 µL) was then transferred into a 5 mm NMR tube for spectral analysis.

NMR spectra were acquired using a proprietary pulse program developed by Maven Health, specifically optimized for saliva metabolomics. All spectra were processed using the software’s automated procedures for baseline correction, phasing, and referencing to the internal TMSP standard.

### Statistical analysis

For statistical analysis, differences between metabolite concentrations across sample groups were assessed using the limma package v3.54.2 (linear models for microarray data) in R. The limma method was employed to fit linear models and compute empirical Bayes statistics for differential abundance testing. Significance was determined using adjusted p-values, and metabolites with a false discovery rate (FDR) below 0.05 were considered significant.

To examine a potential statistically significant difference between the average weight change over time for each participant, a paired samples t-test was conducted. The analysis was performed using the scipy.stats.ttest_rel function in Python (scipy v. 1.9.1).

### Bio-energetic calculations

The gold standard for calculating the resting metabolic rate (RMR) is considered to be indirect calorimetry. However, as these measurements were beyond the scope of this study, we used the Mifflin-St. Jeor equation. This equation is generally considered to be the most accurate of various RMR equations. Any equation, including the Mifflin-St. Jeor, have their limitations^9^ and should be used with caution in stratified groups within a population (by sex, ethnicity, age, weight classification etc.). The Mifflin-St. Jeor equation takes weight, height, age, and the sex of the individual as variables. The equation was established by performing multiple linear regression analyses on 498 healthy subjects, with a group of participants of various ages and obesity levels. The RMR is highlighted in Equation 1.

After calculating the *RMR_person_*, the caloric deficit over the course of the program was calculated. We did not adjust the *RMR_person_* during the program (Even though the weight varies). We simplified the weight variable during the program since the initial weight loss during the program is most likely attributed to, among other factors, a change in fluid retention. We define the caloric deficit in kcal as follows in Equation 1.

We observed (Figure 3) that the measurements of acetone exceeded the reference range maximum values during Day 2 (Reference ranges from Healthy non-fasting cohort as defined by Maven Health research trial (unpublished). This led us to assume that ketogenesis became the main source of energy after Day 1. Therefore, we adapted the caloric deficit estimation, to specifically measure the deficit that is compensated by burning fat.

Once the ketogenic period caloric deficit was known for each participant, we aimed to estimate how much fat (in grams) was used to compensate for this caloric deficit. Various assumptions were made, further explored in Table 4.

**Table 4.** Bio-energetic calculation assumptions with rationale and an effect estimation for each assumption.

| Assumption | Rationale | Over-<br>underestimation |
| --- | --- | --- |
| (body) fat equals white adipose fat tissue (WAT) | WAT accounts for 80-90% of a healthy adult | No effect |
| Fatty acid composition in WAT is consists of Oleic acid, Palmitic acid and Linoleic acid in a 3:2:1 ration respectively (50%, 33.33% and 16.66%). | WAT contains many fatty acids, whereas oleate, palmitate and lineolate represent the bulk (85%) of all triglyceride rich droplets in adipocytes.<sup>10</sup> | No effect |
| Triglycerides consist of 3 fatty acids and 1 glycerol. A Fatty acid produces 114 ATP and glycerol 20 ATP. One triglyceride produces 362 ATP in total. | Oleate (118.5 ATP), palmitate (106 ATP) and lineolate (117 ATP) generate ATP through Beta-oxidation and oxidative phosphorylation. Averaged out in 3:2:1 ratio, the average fatty acid produces 114.08 ATP. We assume 114 ATP per fatty acid. | Increases the general error due to varying fatty acid contents dependent on diet and other factors in each participant. |
| The molecular weight of a typical triglyceride molecule in humans is around 916 g/mol | Oleate - C<sub>18</sub>H<sub>34</sub>O<sub>2</sub> - 282.45 g/mol<br>Palmitate - C<sub>16</sub>H<sub>32</sub>O<sub>2</sub> - 256.42 g/mol<br>Linoleic - C<sub>18</sub>H<sub>32</sub>O<sub>2</sub> - 280.44 g/mol<br>Avg. Mol weight: 274.5 g/mol<br>Glycerol - C<sub>3</sub>H<sub>8</sub>O<sub>3</sub> - 92.09 g/mol<br>Total: 3 x 274.5 = 92.09 = 915.59.<br>Simplified 916 g/mol | No effect |
| To replace 1 kcal deficit, we require 8.22 x 10<sup>22</sup> ATP molecules<sup>11</sup>. | 8.22 x 10<sup>22</sup> molecules of ATP are generated per 1 kcal at a '100%' economy. Mitochondria can operate at different effectiveness levels. We assume a 100% level with the potential to underestimate calorie level as a conservative estimate, as we have not measured the effectiveness of mitochondria in the participants. | Underestimation of the calories needed and thus potential fat burn. |

*Equation 1 Set of resting metabolic rate (RMR), caloric deficit estimation & ketogenic period caloric deficit estimation*

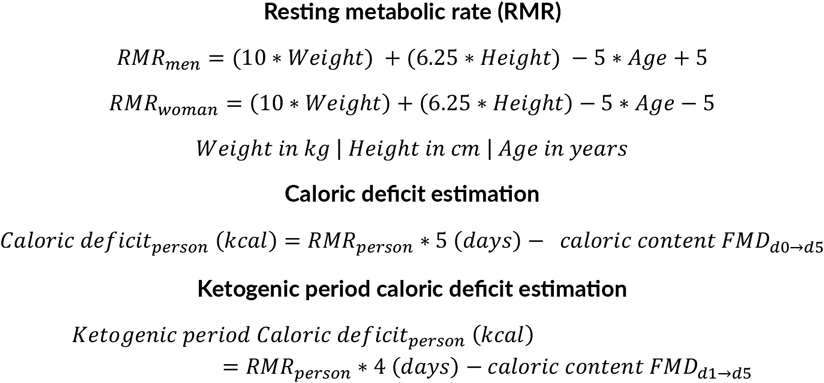

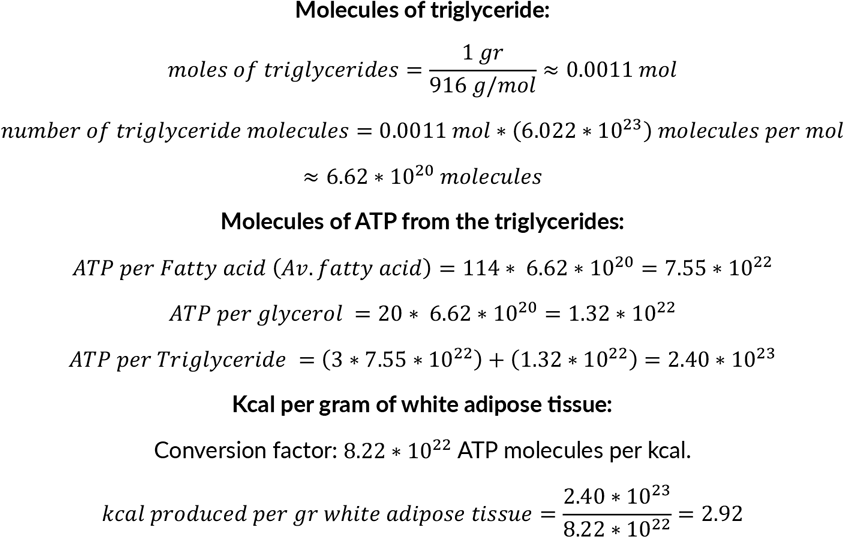

We calculated how much kcal 1gr of white adipose tissue (fat) can generate. By combining the abovementioned assumptions in Table 4 and using Avogadro’s number for g/mol to molecules conversions we can complete the equations.

## Results

In this study 6 participants were subjected to a 5-day reSET Fasting Mimicking Diet (FMD) program. The main characteristics of this cohort are summarized in Table 5. The saliva metabolome showed changes in various metabolic domains. All significant metabolite changes can be viewed in Figure 1. A strong significant increase of the ketogenesis by-product acetone was observed in every participant, alongside significant weight loss right after and up to 3 days after refeeding. Various metabolites serving as energy substrates in mitochondrial energy pathways were seen to be significantly affected, implying energetic reprogramming. Strong effects were also observed in metabolites derived from the microbiome, suggesting enhanced activity.

**Table 5.** Cohort characteristics.

| Participant properties (n=6) | Mean ± SD | Median (IQR) |
| --- | --- | --- |
| Age (years) | 37 ± 5 | 38 (34-42) |
| Height (cm) | 171 ± 10 | 171 (163-178) |
| Weight (kg) | 71 ± 10 | 73.65 (72-76) |
| BMI (kg/m<sup>2</sup>) | 24 ± 4 | 24 (22-26) |

**Figure 1.**
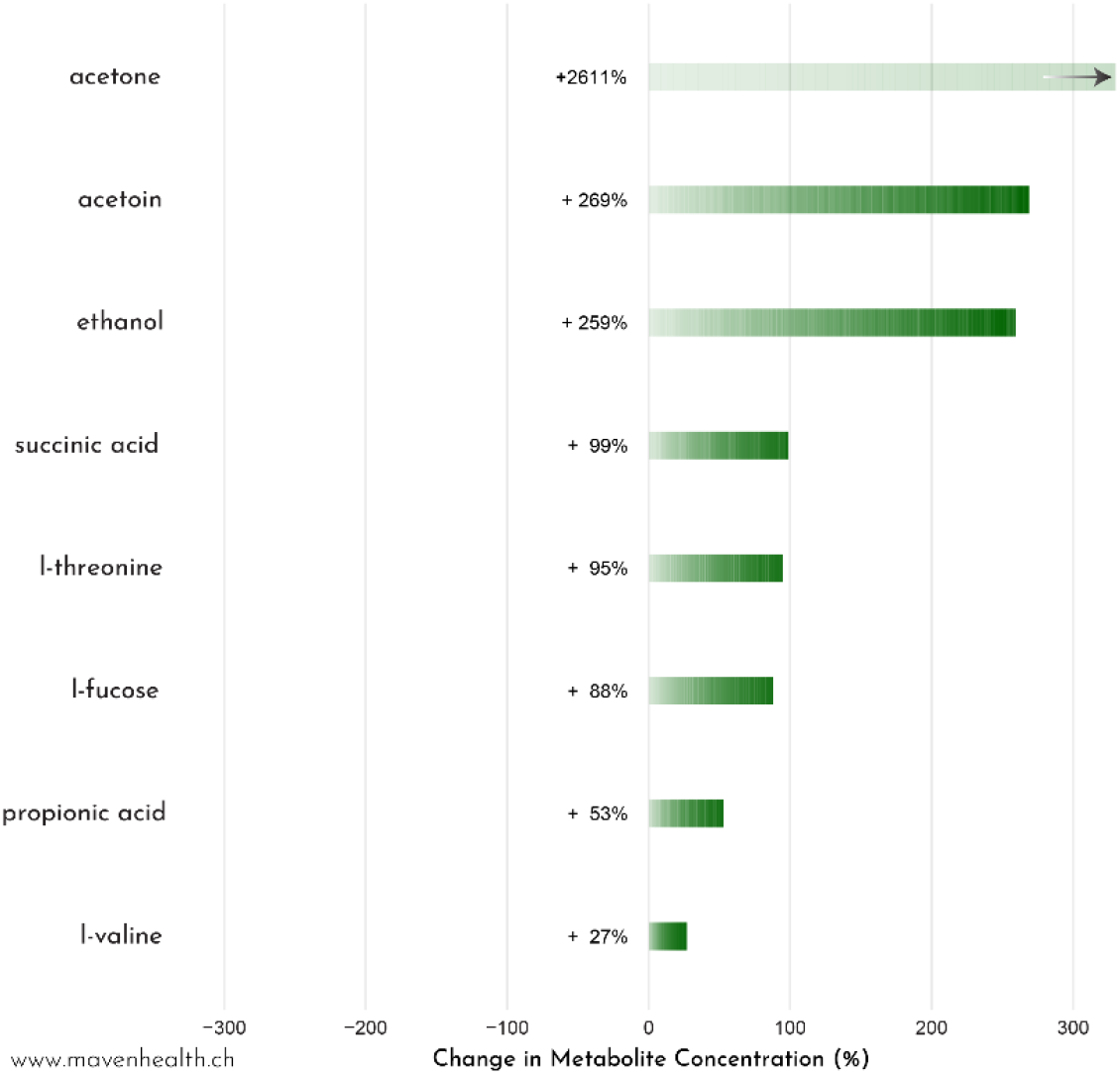
Significantly changed metabolites measured in saliva over the fasting-mimicking diet (N=6, p < 0.05). Green bars represent metabolites with increased levels, with average percentage changes labeled beside each bar. Percentage changes were calculated by comparing baseline values on Day 0 (non-fasting) with final quantities measured at the end of the diet Day 6 (fasting).

### FMD impact on weight loss largely due to fat loss

FMDs are known to lead to weight loss. In a randomized cross-over study^3^ with 100 generally healthy volunteers the FMD was monitored over three months with 5 days of FMD each month. In this study the authors found that the FMD participants lost −2.6 ± 2.5 kg in body weight and reduced the total body fat by mass −1.393 ± 1.786 kg. Thus, an estimated 54% of weight loss can be attributed to fat loss in the case of a 3-month repeated FMD scenario.

In this section we investigate how the human body produces energy during periods of reduced caloric intake. We present a novel method to assess the total body fat loss in comparison to the (sustained) weight loss during and after completion of the program using the self-reported data and the Maven Health Metabolic Insights test.

### Energy generation

During an FMD the body initially uses up the available glucose. This starts with the breakdown of the glycogen stores and subsequently from the de-novo glucose synthesis from non-carbohydrates, notably certain amino acids.

After the glucose stores are depleted, fat breakdown (lipolysis) begins closely followed by ketogenesis and ketolysis. Ketogenesis is the process by which ketone bodies are synthesized from fatty acids and specific ketogenic amino acids. These ketone bodies can subsequently be broken down through the process of ketolysis to produce energy. The ketone bodies produced during ketogenesis are acetoacetate, beta-hydroxybutyrate, and acetone. Whereas the first two will be used during ketolysis to generate energy, the third ketone body, acetone, is a byproduct. Acetone is expelled from the body through exhalation and urination. After extended or prolonged fasting, tissues might be broken down for essential glucose production. In a late starvation phase, once all fat stores are exhausted, protein degradation accelerates which can lead to starvation-related symptoms. A visual representation of these processes can be observed as a time sequence in Figure 2.

**Figure 2.**
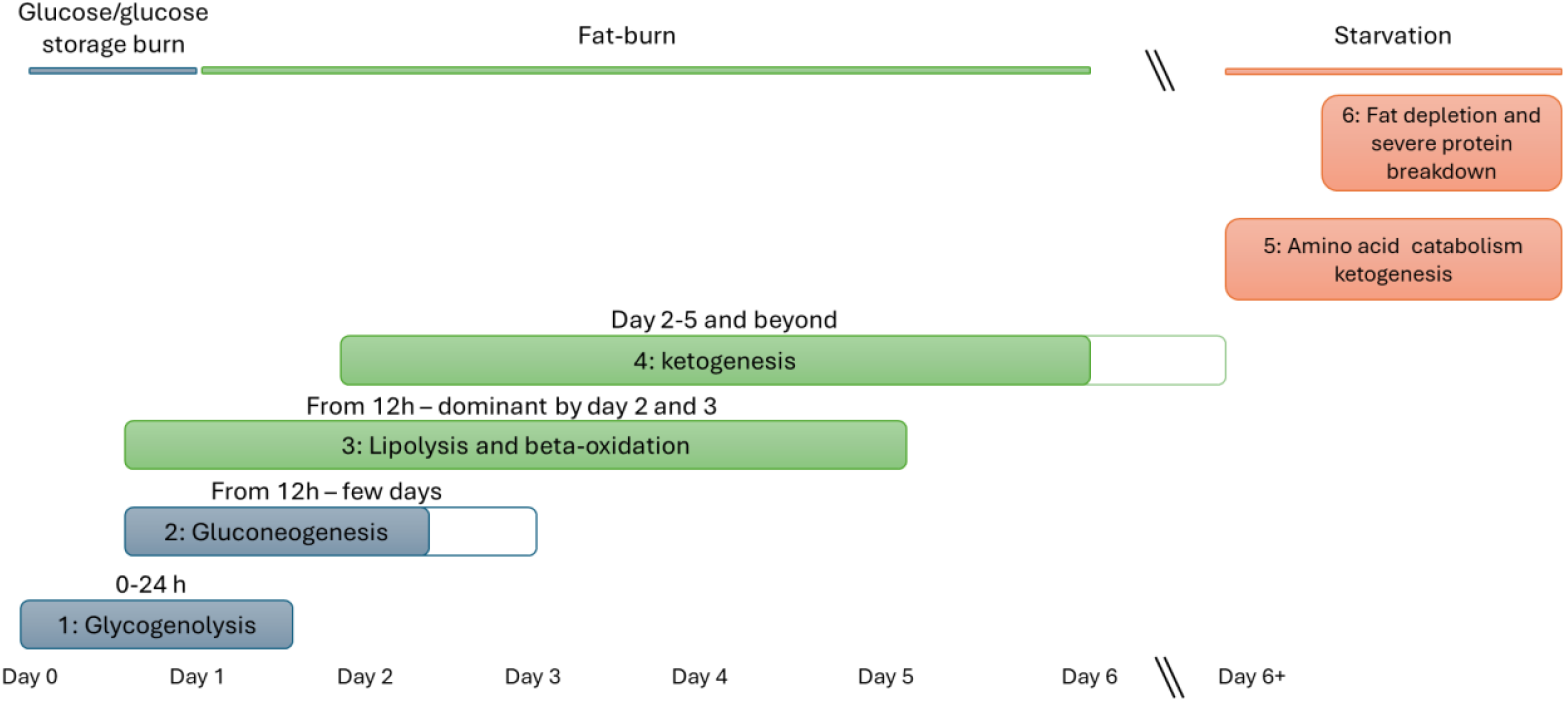
energy utilization in the human body during caloric deficits and eventual starvation.

FMDs are designed to have a person reach the fat burning stage and not to lead to the starvation and muscle-breakdown phase, as these are evidently last resorts for the human body to generate energy. Starvation can have severe consequences and should be avoided^12^.

### Bioenergetic calculations

To perform an accurate estimation of the fat burn in each participant without the use of indirect calorimetry or imaging techniques like Dual-Energy X-ray Absorptiometry (DEXA) we have produced a bioenergetic calculation framework. This framework includes the caloric needs and deficit during the FMD and estimations from when ketogenesis provides the energy required to compensate for said caloric deficit. This framework includes the caloric needs (calculated via the *resting metabolic rate*) and the *active ketogenesis period*.

#### Resting metabolic rate

The resting metabolic rate (RMR) helps us express the amount of energy that an individual requires to keep the body functioning at rest. Due to the caloric restriction of the FMD program, participants reported not to have performed heavy exercise like strength training or cardio. We assumed that light walks and normal daily activities did not meaningfully impact the RMR of the participants during the program. Additionally, lean body mass and fat mass contents could meaningfully impact the RMR. However, these were not considered in our participant’s RMR calculations.

#### Active ketogenesis period

Acetone is produced as a byproduct of ketogenesis and can build up in the body during fasts or periods of reduced carbohydrate intake^13^. The body eliminates acetone through two pathways: 1) respiration; acetone diffuses into the bloodstream and is then expelled through the breath and 2) urination; filtered by the kidneys and excreted in urine.

As shown in Figure 3, ketogenesis marks the beginning of the burning of fat. The byproduct acetone was measured in the saliva with Maven Health’s Metabolic Insights test. Acetone increased in all six participants during the FMD program. Notably during day 2 the levels exceeded the maximum of the biological range measured in Maven Health’s completed healthy cohort study (unpublished).

**Figure 3.**
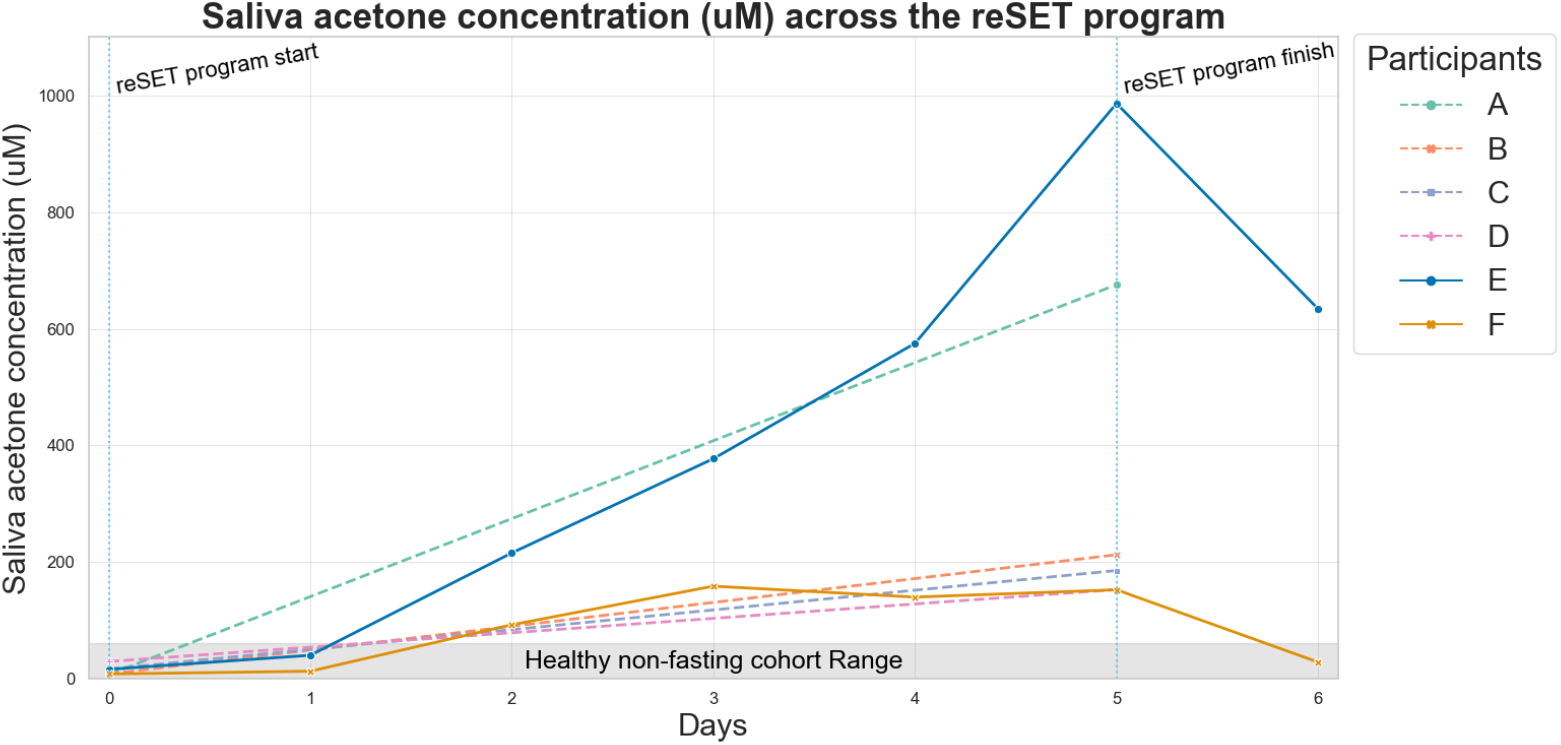
Absolute acetone levels (uM) over time for all participants. (Participant A-D have a t0 and t5 measurement; Participant E-F have daily measurements from t0 – t6); Healthy non-fasting cohort Range as defined by Maven Health research trial (unpublished; range 1.78 - 60.58 uM).

Based on these results we estimated that ketogenesis started to be the main method of generating energy starting from day 2. For our bioenergetic calculations we assumed that during day 1, the first day of dieting, no fat is burned, and starting day 2 ketogenesis starts and is responsible for the full caloric deficit energy requirements.

Using the RMR together with the daily caloric intake we establish the caloric deficit for the days of the program where ketogenesis is used to compensate for the energy deficit.

We calculated the amount of white adipose tissue in grams is required to be burned to compensate for the ketogenic period caloric deficit. Core assumptions were as follows: 1) White adipose tissue is exclusively used as a fat source.

2) The white adipose tissue’s fatty acids consist solely of oleate, palmitate and lineolate in equal fractions (as they represent the bulk (85%) of all triglyceride rich droplets in adipocytes^10^). 3) We did not adjust the RMR daily and use the day 0 RMR throughout the calculations.

### Weight

Significant weight loss was observed in the cohort from the start to finish of the program in all participants and within the male and female subgroups. Sustained weight loss was observed from the start of the program up to 3 days after refeeding (day 8). The results are summarized in Table 6. The progression of weight over time can be seen in Figure 4. A rebound effect can be seen on Day 6. This is most likely due to the fact that participants returned to their standard diet, impacting the fluid retention. The observed weight loss was consistent for all participants. A normalized perspective of the timeline of the weight change can be seen in Figure 5.

**Table 6.** Weight change during the FMD program, rebound effect (day5->day8) and the full program and refeeding effect (d0->d8)

|  | All (n=6) | Male (n=3) | Female (n=3) |
| --- | --- | --- | --- |
| Weight change<sub>d0->d5</sub> (kg) | -2.84*** | -3.07* | -2.62* |
| Weight change<sub>d5->d8</sub> (kg) | 0.62* | 0.7 | 0.55 |
| Weight change<sub>d0->d8</sub> (kg) | -2.22** | -2.37 | -2.07 |

(paired t-test; p-values: \* < 0.05; \*\* < 0.01; \*\*\* < 0.001)

**Figure 4.**
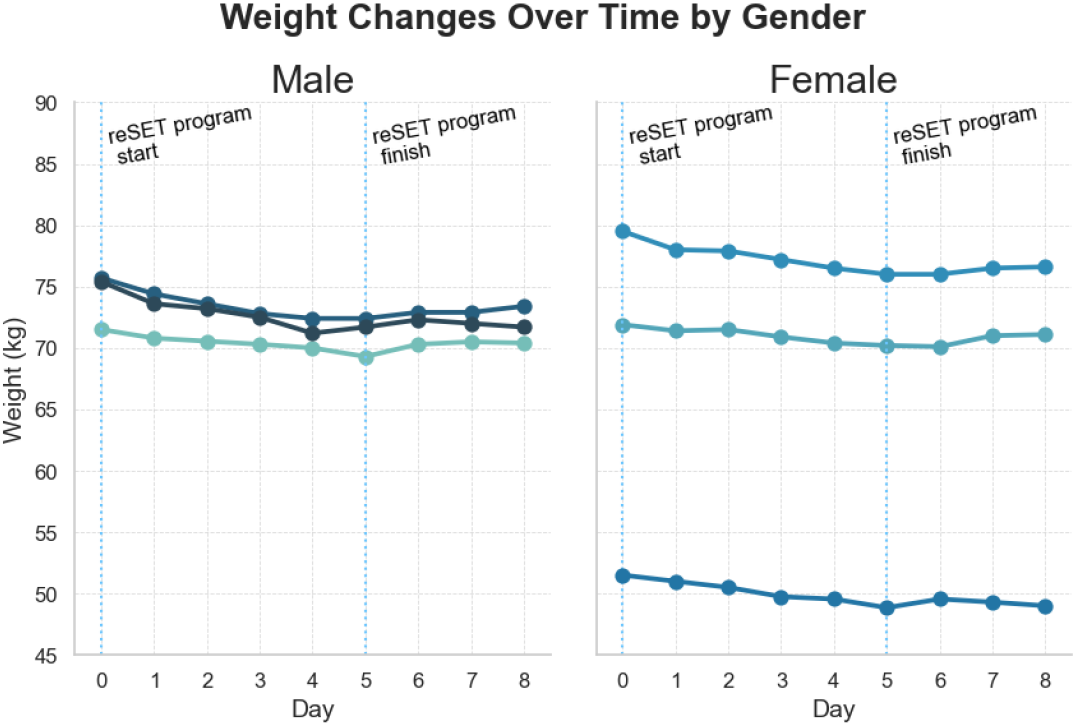
Weight (kg) progression over the full duration of the program and up to 3 days after program finish.

**Figure 5.**
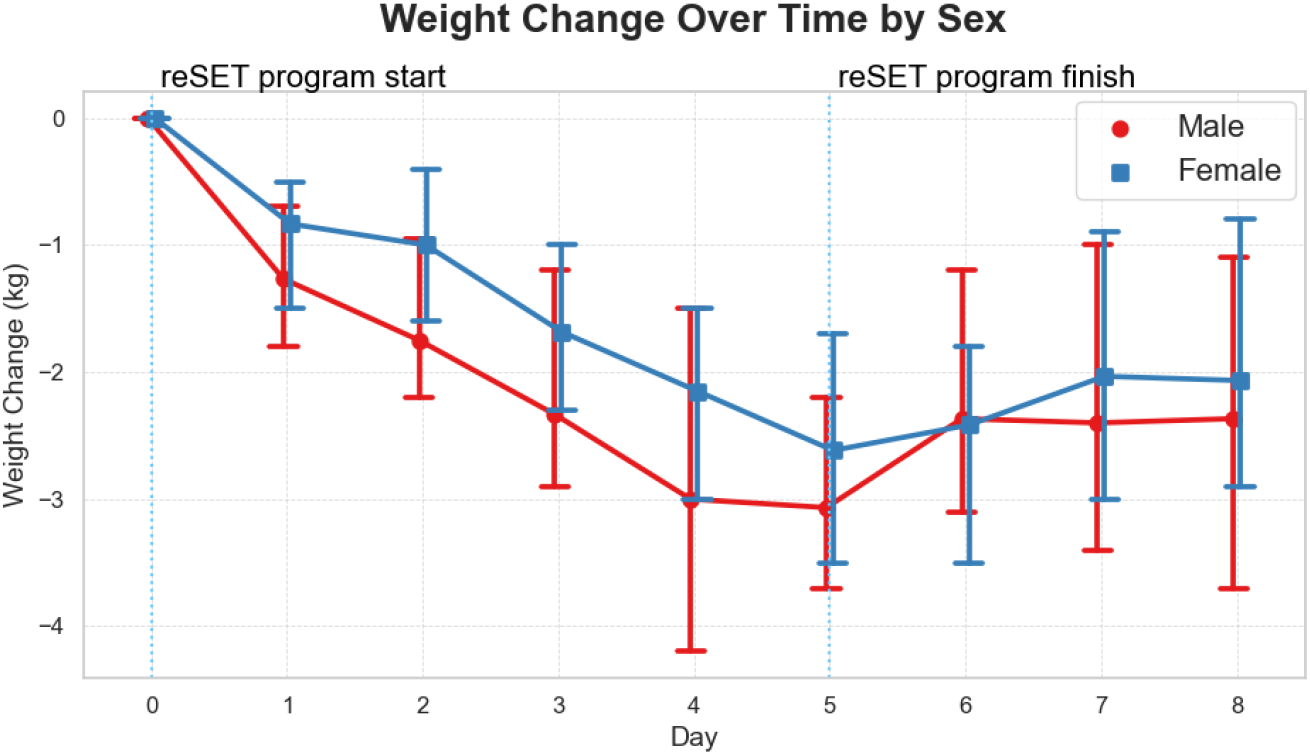
Normalized weight loss (kg) over time by Sex. Day 0 (non-fasting), Day 5 (fasting; end) and day 8 (non-fasting; 3 days of refeeding/return to normal diet).

### Fat burn

The observed average weight loss could be driven by several factors. To assess to which degree the weight loss can be attributed to fat loss, we performed bio-energetic calculations to estimate how much fat was burned during the program. Briefly, we used the Mifflin St-Jeor equation (Equation 1) to estimate the caloric deficit per participant. We then converted this energy deficit to kg fat tissue following the assumptions in table 4. Our findings are summarized in table 7. We found that, on average, a participant reaches a cumulative caloric deficit of 2924 kcal over the ketogenic period of 5 days fasting programme which correlates to roughly 1 kg of fat burned to maintain metabolic homeostasis. Thus, considering a total weight loss of 2.22 kg we estimate the fat loss to be roughly 45% of the total sustained weight loss.

**Table 7.** Estimated fat burned, RMR and total caloric deficit during the FMD program.

|  | All (n=6) | Men (n=3) | Women (n=3) |
| --- | --- | --- | --- |
| Potential fat burned (kg) | -1.01 ± 0.27 | -1.22 ± 0.98 | -0.78 ± 0.20 |
| Resting metabolic rate (kcal) | 1513 ± 199 | 1676 ± 70 | 1351 ± 147 |
| Total Caloric deficit<sub>d1->d6</sub> (kcal) | -2924 ± 796 | -3572 ± 287 | -2277 ± 589 |

(mean ± standard deviation)

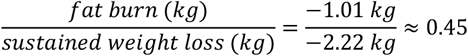

Concretely we have observed that in men, the fat burn during the program could be as high as 1’092–1’328 grams and in women 497–952 grams. During the last day (d_4->5_) of the program, where the caloric deficit is the highest, the fat burn could be as high as 281-339 gr and 131-245 gr in men and women respectively.

### FMD increases mitochondrial metabolites and induces Energetic Reprogramming

During periods of metabolic excess, the human body is in a persistent anabolic state. This is paired with failure to activate lysosomal and autophagy states, leading to fibrosis, excess proliferation, and changes in immunometabolic phenotype^14^. These effects contribute to chronic non-communicable diseases such as diabetes and heart attacks. Cellular metabolism in a persistent anabolic state mostly relies on aerobic glycolysis, named the Warburg effect. Fasting or exercise have profound beneficial effects on the human metabolism that counteract these detrimental processes. During periods of exercise or fasting, humans enter a so-called “anti-Warburg effect” state where, due to the higher demand versus supply, the generation of energy shifts to a more efficient state of mitochondrial respiration.

In this study, we observed that the FMD cohort showed metabolic shifts that align with energetic reprogramming and the “anti-Warburg effect”. We compared key metabolites involved in the TCA cycle between the non-fasting and fasting groups. Besides the shift to a ketone-based metabolism, we observed significant increases in key metabolites involved in the TCA cycle: L-glutamine, L-glutamate and succinic acid (Figure 6). These metabolites are key substrates for the TCA cycle and reflect energetic metabolic reprogramming towards enhanced mitochondrial function and energy efficiency (Figure 7).

**Figure 6.**
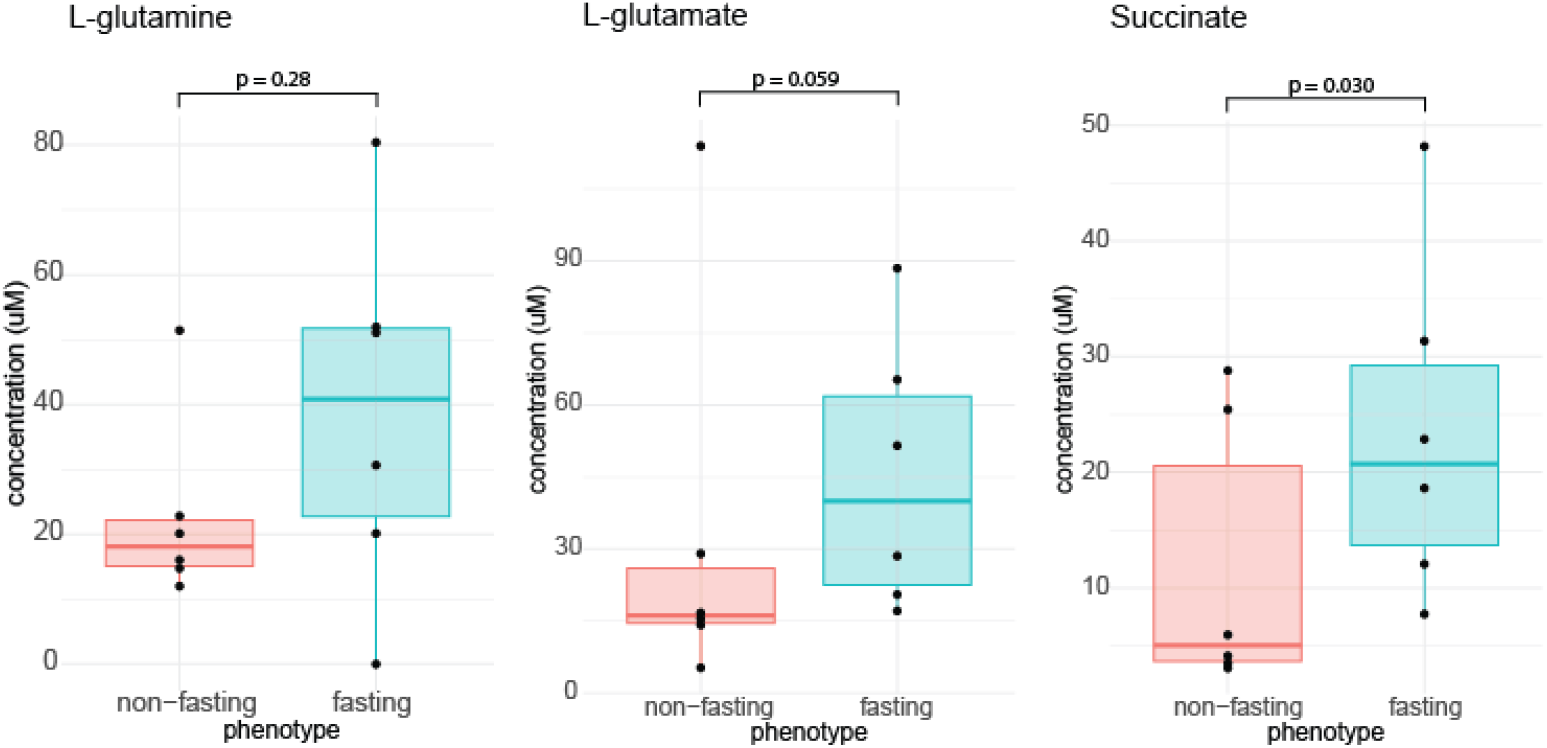
Boxplots showing the concentrations of metabolites involved in the TCA cycle. Individual measurements are indicated by black dots. P values of the differences are indicated above each plot. Left boxplot (red): non-fasted baseline measurements. Right boxplot (blue): fasting measurement, taken on day 5. N = 6 paired biological measurements for each group.

**Figure 7.**
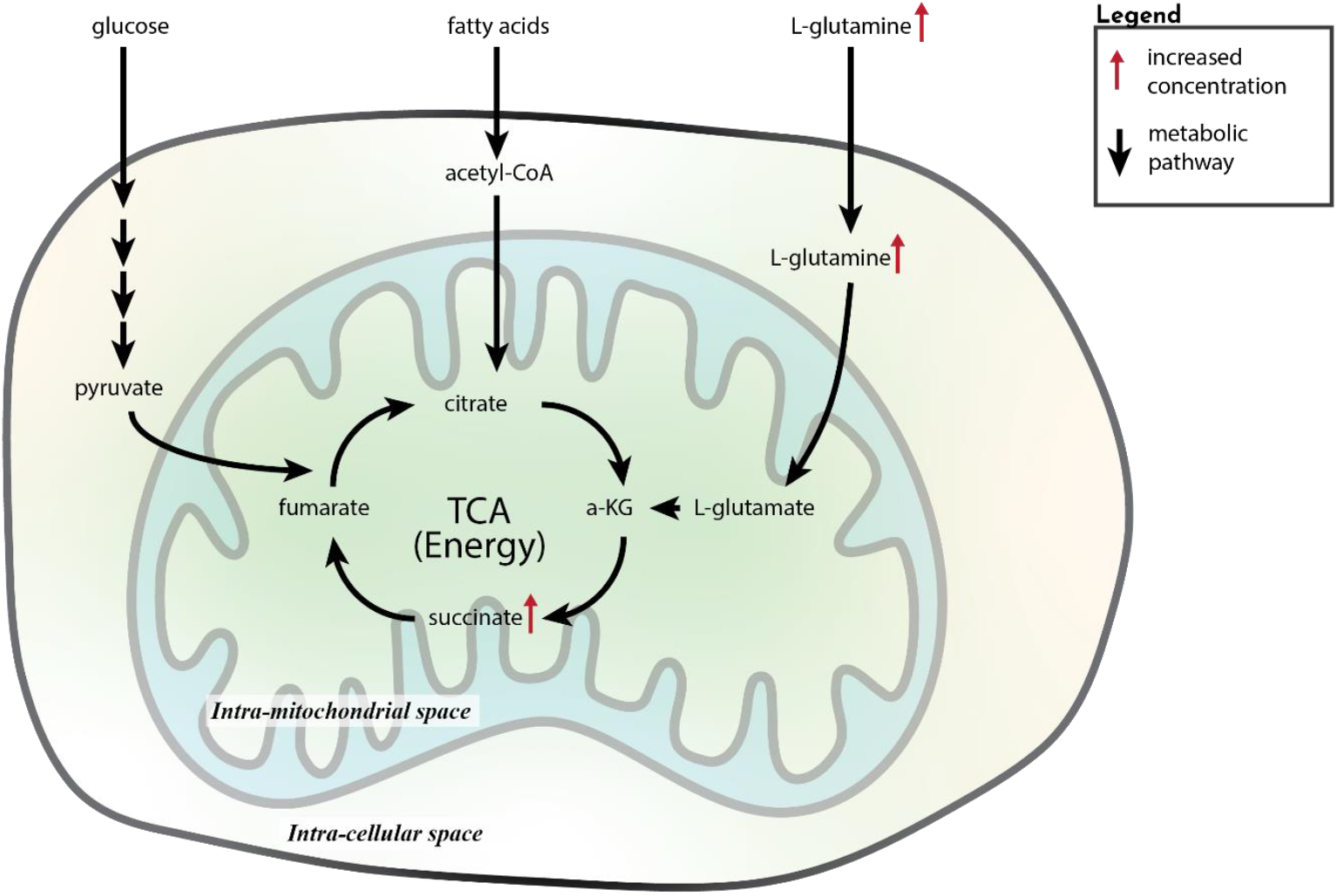
Schematic illustrating the flow of substrates and associated pathways in the mitochondrion. This schematic represents the integration of multiple metabolic pathways that contribute to the tricarboxylic acid (TCA) cycle, highlighting energy production and key metabolite exchanges between the intra-cellular space and the intra-mitochondrial space. Red arrows indicated increased concentrations of metabolites between the two measurement days. Black arrows indicated the metabolic pathways. Multiple arrows indicate a number of intermediate steps.

### FMD stimulates microbiome metabolic activity

The human microbiome plays an important role in not only host defense and digestion. Metabolites produced by the human microbiome also play a major role in the proper functioning of several metabolic processes, the main metabolite effectors being the short-chain fatty acids. In this study we investigated if the FMD diet improved the microbiome metabolism. We compared the levels of substrates and microbiome derived metabolites between the non-fasted and fasted states.

A marked increase in the levels of the short-chain fatty acids propionic acid, butyric acid, and acetic acid was observed between the initial non-fasted state and the post-FMD state (Figure 8). In addition, we observed the increases in L-fucose, a crucial metabolite produced by the gut microbiota which plays a role in gut health and immune function. Additionally, we observed increases in acetoin and ethanol, which are produced by fermentation of the provided carbohydrates, indicating an active microbial metabolism that promotes a healthy gut environment.

**Figure 8.**
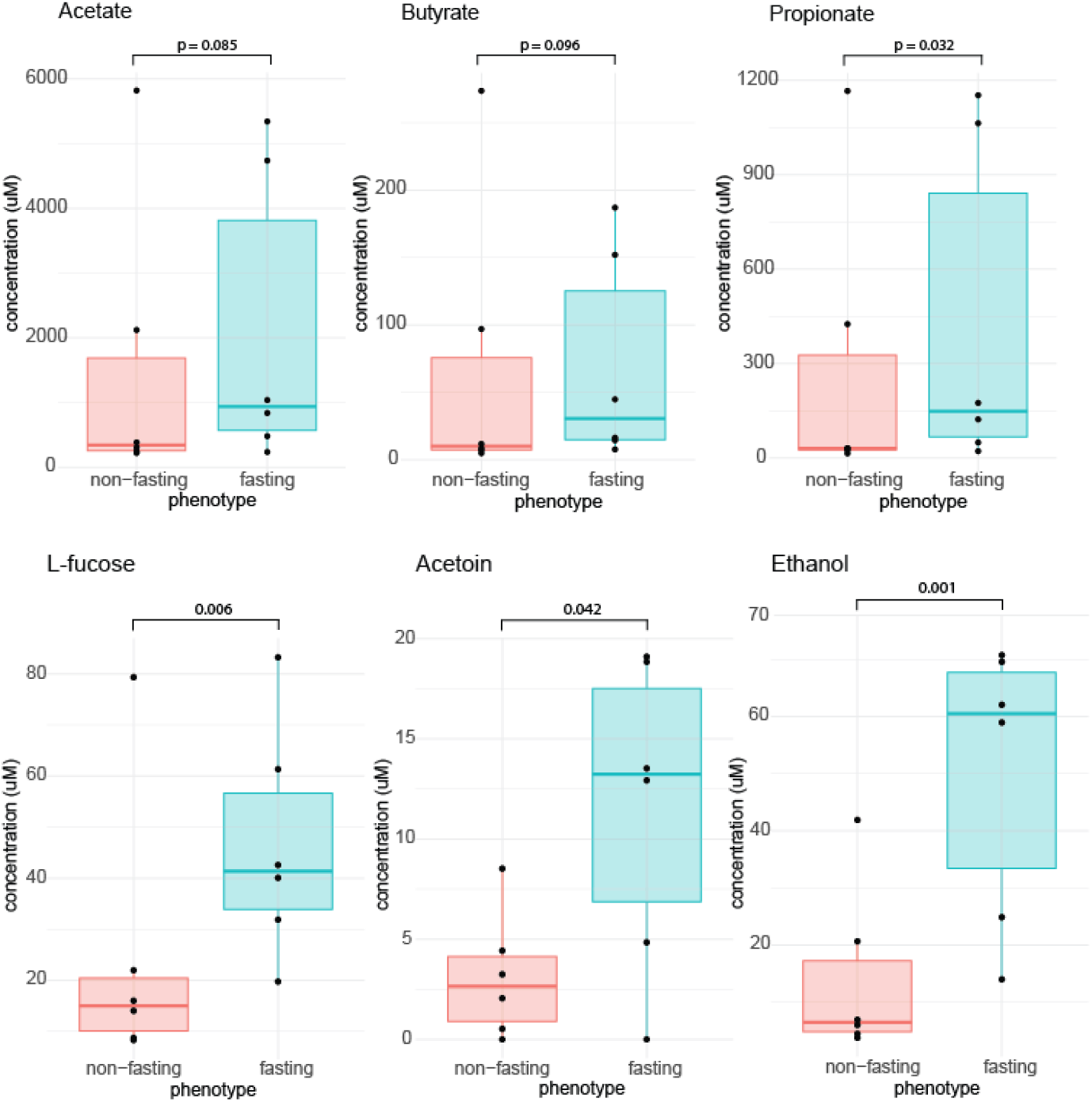
Boxplots showing the concentrations of Short-Chain Fatty Acids (SCFAs) and other various metabolites as indicated, produced by the gut microbiome. Individual measurements are indicated by black dots. P values of the differences are indicated above each plot. Left boxplot (red): non-fasted baseline measurements. Right boxplot (blue): fasting measurement, taken on last day of program. N = 6 paired biological measurements for each group.

## Discussion

This study found notable metabolic effects of the FMD in the participants and should be regarded as an initial exploration into how fasting and FMDs impact human physiology. Due to the limited sample size (n=6) of the study, it was not expected to find statistically significant effects. However, multiple statistically significant effects were observed (Figure 1) indicating that the FMD used in this study most likely has a powerful effect that can be replicated in larger cohorts across multiple populations subgroups. The ability to perform daily testing during studies or routine client monitoring greatly enhanced the ease to perform the testing and the collection of the samples. In fact, a common limitation in many studies is often the need of participants to visit a clinic or research facility for each test moment. With an easy-to-use non-invasive saliva test testing can be done with convenience and at a flexible location (i.e. the person’s home). In combination with not requiring training for the collection creates huge potential value for future intervention studies.

Non-invasive and cheap methods that can assess the body fat loss in fasting participants are of interest as the weight loss during and after FMDs does not equal total body fat loss. Multiple factors play a role in the total composition of weight loss and not all of them are lasting effects. Notably a rebalancing of the fluid retention in the human body. The change to a low-salt FMD may lead to a tendency for lower fluid retention during the first days until the body reaches a new fluid homeostasis.

The fat loss as a component of total weight loss estimated by bio-energetic calculations, resulted in biologically relevant values. However, no validation by DEXA or indirect calorimetry was performed to verify the calculated fat loss result. Therefore, the bio-energetic framework is a theoretical work and must be verified by other methods. As the energy metabolism is complex, many assumptions were made with potential over- and underestimations (highlighted in the material and methods). A cumulative error of the various assumptions might lead to a sizable overall error, thus practical use of the approach should be used with caution.

FMDs are severely calory-restricted, with a severe limitation on carbohydrates as an energy source. During calory-restricted diets such as the one studied here, mitochondria increase their bioenergetic capacity to compensate for the reduced dietary availability of energy through remodeling, reshaping, and increased activity of the respiratory chain complexes. Prompted by the lack of glucose as an energy source, mitochondria react by adapting to alternative energy sources, the primary being fatty acids and ketone bodies, liberated through the lipolysis of triacylglycerols stored in adipose tissue. These metabolic pathways converge on the main energy carrier molecule acetyl-CoA, which is metabolized in the TCA-cycle. Besides using acetyl-CoA as fuel for the TCA-cycle, the mitochondria increase the influx of other TCA-cycle intermediates, notably through the L-glutamate, L-glutamine, and succinate axis. L-glutamine is a key substrate for mitochondrial energy production. L-glutamine is converted into L-glutamate which enters the TCA cycle via conversion to alpha-ketoglutarate, further supporting energy generation in mitochondria. Succinic acid is then produced from alpha-ketoglutarate in the TCA cycle, and its rise reflects increased mitochondrial activity, enhancing ATP production through oxidative metabolism.

In this study we observed measurable increases in the key TCA-substrates and precursors L-glutamine, L-glutamate, and succinate. As shown in Figure 8, we saw marked increases in all measured metabolites of this metabolic axis. Changes in metabolites were measured before and after the FMD, which demonstrates the energetic remodeling of mitochondrial and improved bioenergetic capacity. These findings suggest that the FMD actively reduces the reliance on glycolysis as a source of ATP, providing tangible health benefits such as reduced immune-inflammatory phenotype and lower risk of chronic non-communicable diseases^14^. As described in the previous section, levels of acetone - the main byproduct of fat metabolism and ketogenesis - increased, demonstrating a shift to lipid metabolism. This metabolic shift from glucose to fat-derived energy sources further reduces the dependency on glycolysis.

The human digestive tract is host to a diverse array of microorganisms, including bacteria, fungi and protozoa. The bacteria in humans’ digestive tract play an important role not only in digestion but have strong physiological effects on our health. A healthy microbiome has a profound positive impact on behavior, immunological regulation, energy homeostasis and metabolism^15^. The signaling molecules through which the microbiome elicits these health benefits are the short-chain fatty acids (SCFAs): acetate, butyrate, and propionate^16^. SCFAs help maintain epithelial barrier integrity in the gut by promoting mucin expression, tight junction formation, and antimicrobial peptide production. Specifically, butyrate and propionate exert strong anti-inflammatory effects, whereas propionate and acetate are involved in fatty acid oxidation and influence metabolic pathways.

The FMD used in the ‘reSET’ program is based on the work of Dr. Valter Longo^17,18^. The focus is a high fat content from healthy sources, such as olive oil, nuts, and seeds. This high fat content supports microbial diversity and provides a rich substrate for fermentation. The rationale is that the growth of beneficial gut bacteria is promoted and the production of SCFAs is enhanced. Furthermore, the inclusion of whole grains, fruits, and vegetables in addition to healthy coconut-based bites provide prebiotic fibers that further nourish beneficial gut flora. In this study we observed a marked increase in metabolites originating from the microbiome. All three measured short-chain fatty acids show an upward trend after the five-day fasting trial. Besides these key molecules, the microbiome also produces other relevant metabolites such as acetoin, L-fucose and ethanol. The diet was rich in fibers, with a daily snack consisting mostly of dried fruit and coconut fibers, providing a rich source of substrates for the gut microbiota. The increase of microbiome metabolic health has been shown to have numerous health benefits for the host^16,19^. L-fucose is a sugar which is produced by humans and is used by bacteria to produce the short-chain fatty acids mentioned above. However, L-fucose is also the fundamental sub-unit of the polysaccharide fucoidan, found in the cell wall of seaweed and algae. The diets included a daily intake of 6 spirulina tablets, composed of a type of blue-green algae. It is thus unclear to what degree the increase in L-fucose stems from endogenous production of from dietary supplementation.

## Conclusion

This study provides valuable insights into the effects of fasting and FMDs. The findings indicate that FMDs can promote weight loss, alter bioenergetic substrate utilization, and enhance gut microbiome activity. Given the small sample size (n=6), the observed significant effects suggest substantial biological changes, though these results should be interpreted as exploratory. Further research with larger, more diverse cohorts is essential to confirm and expand upon these initial observations.

Our findings suggest that the reSET diet has potential as an effective approach for inducing weight loss and fat loss. The FMD alters energy production pathways in the body, which may be associated with numerous health benefits documented in existing literature. Increased gut microbiome activity was also observed, a change often linked to improved health outcomes.

The testing of saliva metabolites yielded a rich dataset, demonstrating this method’s feasibility as a simple, non-invasive approach to monitor the physiological impacts of lifestyle and dietary interventions. Saliva testing offers significant potential for supporting frequent, user-friendly testing in future studies and real-world applications.

## Disclaimer

This white paper was written as a result of a product collaboration between Eat By Alex and Maven Health. All research design, data analysis, and interpretation of results were conducted independently by Maven Health. The conclusions and recommendations presented here are based solely on Maven Health’s objective analysis of the data. Eat By Alex provided compensation for running the analysis of the saliva samples to Maven Health and provided the ‘reSET’ diet to the participants at no cost. The participants consented to participate in this product collaboration and are employees from Eat By Alex (n=4) and Maven Health (n=2). The goal of this collaboration is to leverage the synergy of both companies’ approach to offer an improved product to their customers.

### Maven Health GmbH

Maven Health is a Zurich-based company that offers healthcare professionals, like dieticians, nutritionists and doctors a non-invasive saliva testing service that analyses key metabolites and compares the end-customers results with cohorts of healthy individuals’ saliva values. Maven Health’s testing results are delivered in intuitive reports with personalized scores, tailored metabolite insights and nutritional recommendations.

### Eat By Alex AG

Eat by Alex is a Zürich-based wellness company offering holistic, science-based health programs to empower individuals on their journey to well-being. Through expertly crafted meal plans and comprehensive lifestyle support, Eat by Alex provides solutions for fasting, gut health, stress management, and balanced living, helping clients achieve sustainable health goals.

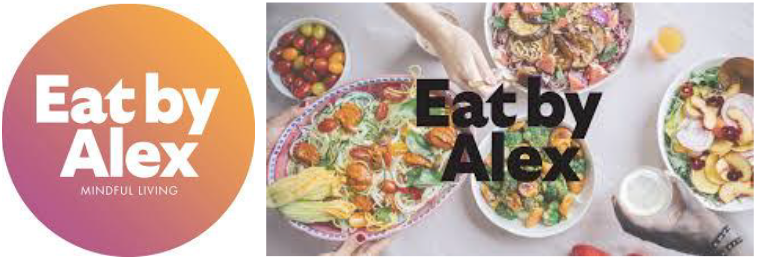

## References

1. Johnstone, A. M. Fasting - The ultimate diet? Obesity Reviews vol. 8 211–222 Preprint at 10.1111/j.1467-789X.2006.00266.x (2007).

2. Brandhorst, S. et al. A Periodic Diet that Mimics Fasting Promotes Multi-System Regeneration, Enhanced Cognitive Performance, and Healthspan. Cell Metab 22, 86–99 (2015).

3. Wei, M. et al. Fasting-Mimicking Diet and Markers/Risk Factors for Aging, Diabetes, Cancer, and Cardiovascular Disease. http://stm.sciencemag.org/.

4. Kenyon, C. The first long-lived mutants: Discovery of the insulin/IGF-1 pathway for ageing. Philosophical Transactions of the Royal Society B: Biological Sciences vol. 366 9–16 Preprint at 10.1098/rstb.2010.0276 (2011).

5. Brown-Borg, H. M. Hormonal control of aging in rodents: The somatotropic axis. Mol Cell Endocrinol 299, 64–71 (2009).

6. Rangan, P. et al. Fasting-Mimicking Diet Modulates Microbiota and Promotes Intestinal Regeneration to Reduce Inflammatory Bowel Disease Pathology. Cell Rep 26, 2704-2719.e6 (2019).

7. Charkey, L.W., Kano, A. K. & Hougham, D. F. Effects of fasting on blood non-protein amino acids in humans. J Nutr 55, 469–80 (1955).

8. Hyvärinen, E., Savolainen, M., Mikkonen, J. J. W. & Kullaa, A. M. Salivary metabolomics for diagnosis and monitoring diseases: Challenges and possibilities. Metabolites vol. 11 Preprint at 10.3390/metabo11090587 (2021).

9. Hasson, R. E., Howe, C. A., Jones, B. L. & Freedson, P. S. Accuracy of four resting metabolic rate prediction equations: Effects of sex, body mass index, age, and race/ethnicity. J Sci Med Sport 14, 344–351 (2011).

10. Yew Tan, C. et al. Adipose tissue fatty acid chain length and mono-unsaturation increases with obesity and insulin resistance. Sci Rep 5, (2015).

11. Ballinger, S. W. Beyond retrograde and anterograde signalling: Mitochondrial-nuclear interactions as a means for evolutionary adaptation and contemporary disease susceptibility. in Biochemical Society Transactions vol. 41 111–117 (2013).

12. Elia, M. Hunger Disease. Clinical Nutrition vol. 19 379–386 Preprint at 10.1054/clnu.2000.0157 (2000).

13. Kalapos, M. P. On the mammalian acetone metabolism: From chemistry to clinical implications. Biochimica et Biophysica Acta - General Subjects vol. 1621 122–139 Preprint at 10.1016/S0304-4165(03)00051-5 (2003).

14. Casanova, A., Wevers, A., Navarro-Ledesma, S. & Pruimboom, L. Mitochondria: It is all about energy. Front Physiol 14, (2023).

15. Sekirov, I., Russell, S. L., Antunes, L. C. M. & Finlay, B. B. Gut microbiota in health and disease. Physiol Rev 90, 859–904 (2010).

16. Mann, E. R., Lam, Y. K. & Uhlig, H. H. Short-chain fatty acids: linking diet, the microbiome and immunity. Nature Reviews Immunology vol. 24 577–595 Preprint at 10.1038/s41577-024-01014-8 (2024).

17. Brandhorst, S. & Longo, V. D. Protein Quantity and Source, Fasting-Mimicking Diets, and Longevity. Advances in Nutrition vol. 10 S340–S350 Preprint at 10.1093/advances/nmz079 (2019).

18. Longo, V. D. & Mattson, M. P. Fasting: Molecular mechanisms and clinical applications. Cell Metabolism vol. 19 181–192 Preprint at 10.1016/j.cmet.2013.12.008 (2014).

19. Kim, J., Jin, Y. S. & Kim, K. H. L-Fucose is involved in human–gut microbiome interactions. Applied Microbiology and Biotechnology vol. 107 3869–3875 Preprint at 10.1007/s00253-023-12527-y (2023).

